# Transcriptome Network Biology Reveals the Neuroprotective Potential of Rosmarinic Acid Against Frontotemporal Dementia

**DOI:** 10.1101/2025.09.05.674389

**Authors:** Amaan Arif, Prekshi Garg, Prachi Srivastava

## Abstract

Frontotemporal dementia (FTD) is a neurodegenerative disorder characterised by impaired behaviour and language. It affects individuals between 45 and 65 years of age. The present study aims to identify potential phytochemicals against target receptor of FTD. Transcriptome data analysis guided computer-aided drug design (CADD) was conducted to investigate molecular interactions between active phytochemicals present in Indian spices and differentially expressed genes. In present study, samples for patients with FTD were obtained from GEO dataset [GSE92340]. The samples were pre-processed followed by differential gene expression analysis using DESeq2. The patient sample was analyzed for 1882 upregulated genes. Hub gene out of DEGs was identified using CytoHubba Plugin of Cytoscape based on eleven parameters (Degree, Edge Percolated Component (EPC), Maximum Neighbourhood Component (MNC), Density of Maximum Neighbourhood Component (DMNC), Maximal Clique Centrality (MCC), Bottleneck, EcCentricity, Closeness, Radiality, Betweenness, and Stress). HDAC1 gene was identified as hub gene and key target in case of FTD. CADD pipeline was used to identify phytochemicals against HDAC1 gene. A list of eighty-one phytochemicals found in Indian spices was analyzed for their drug likeness abilities using molinspiration and toxicity prediction using ProTox-II, which identified twenty-three phytochemicals as safe drug-like compounds. PatchDock was employed to examine interactions between phytochemical compounds and HDAC1. Molecular interactions revealed rosmarinic acid derived from rosemary (Salvia rosmarinus) as the most effective drug for FTD having lowest ACE value of −395 kcal/mol. In conclusion, this study demonstrates essential role of HDAC1 in pathogenesis of FTD and demonstrates the neuroprotective potential of rosmarinic acid against FTD.

## Introduction

Frontotemporal dementia (FTD) is a type of neurodegenerative disorder that affects the frontal and temporal lobes of the brain. The disease is primarily characterized by significant changes in behaviour, language, and sometimes motor function. FTD is generally diagnosed in individuals under 65 years of age, making it a generic form of early-onset dementia.[1] The symptoms of FTD can vary from person to person, but some of the most common symptoms include behavioural changes like apathy, disinhibition, and loss of empathy. Individuals with FTD may also experience language difficulties such as speech problems and word-finding issues. In some cases, motor symptoms like muscle weakness and difficulty with coordination may also be present. [2]

It is important to note that FTD can be challenging to diagnose, as its symptoms often overlap with other neurological disorders. If you or someone you know is experiencing any of the above symptoms, it is recommended to consult with a healthcare professional for a proper evaluation and diagnosis. Frontotemporal dementia is a neurodegenerative disease that currently has no cure. However, treatment options are available to manage symptoms and improve the patient’s quality of life. The treatment approach involves a multidisciplinary team of healthcare professionals who specialize in providing medications, therapies, and supportive care. [3]

Supportive care is an essential part of the treatment plan and includes physical exercises, music therapies, and other activities that can assist in communication and reducing difficulties. Occupational therapy is another vital component of supportive care that can help patients maintain their independence in their daily activities. Medications such as selective serotonin reuptake inhibitors (SSRIs), may be prescribed to control behavioural symptoms like loss of inhibition, overeating and compulsive behaviour. These medications work by increasing the availability of serotonin in the brain, which has a calming effect on the patient’s behaviour. In rare instances, antipsychotics may be used to manage extremely challenging behaviour that poses a risk to patients or others. [4]

It is important to note that treatment plans may vary depending on the patient’s individual needs and response to therapy. Therefore, it is crucial to consult with a healthcare professional to determine the best course of treatment and care plan for each individual patient. Biomarkers are critical in diagnosis of frontotemporal dementia (FTD) and in developing treatment strategies. These are measurable indicators found in biological fluids such as cerebrospinal fluid (CSF) or blood that reflect normal or pathological processes in the brain. FTD biomarkers are used to differentiate it from other types of dementia and track disease progression. Neurofilament light chain, tau protein and TAR-DNA binding protein-43 (TDP-43) are some of the biomarkers associated with specific molecular changes in FTD. By measuring these, researchers gain insights into the underlying mechanisms of the disease and can tailor treatment plans accordingly. This study aims to explore these crucial biomarkers associated with frontotemporal dementia (FTD) by utilizing a network analysis approach. The research begins by using Galaxy Server Europe to identify differentially expressed genes (DEGs) that show significant changes in frontotemporal dementia as compared to healthy individuals (control). [5] These DEGs play significant role in understanding pathological mechanisms of FTD. Cytoscape [6] software analyses DEGs and constructs a network, identifying potential targets for FTD. String Database [7] was used to understand hub genes like HDAC1. CADD pipeline to identify phytochemical compounds could target the HDAC1 gene through Molinspiration [8] and ProToxII [9], and PatchDock [10] to assess interactions with HDAC1. Rosmarinic acid showed strong promise as a treatment option for FTD.

This research aims to provide comprehensive overview of the molecular landscape of FTD through an integrated approach involving DEG analysis, network analysis, functional annotation, and innovative phytochemical screening. The discovery of Rosmarinic acid as a potential therapeutic gives new dimension to interest in effective treatments for FTD, providing hope for individuals who are affected by this condition.

## Materials & Methods

Extensive literature research was conducted to develop a modern Computational Aided Drug Design (CADD) pipeline for this distinct and innovative investigation. The comprehensive pipeline is depicted in Figure 1.

**Figure 1:**
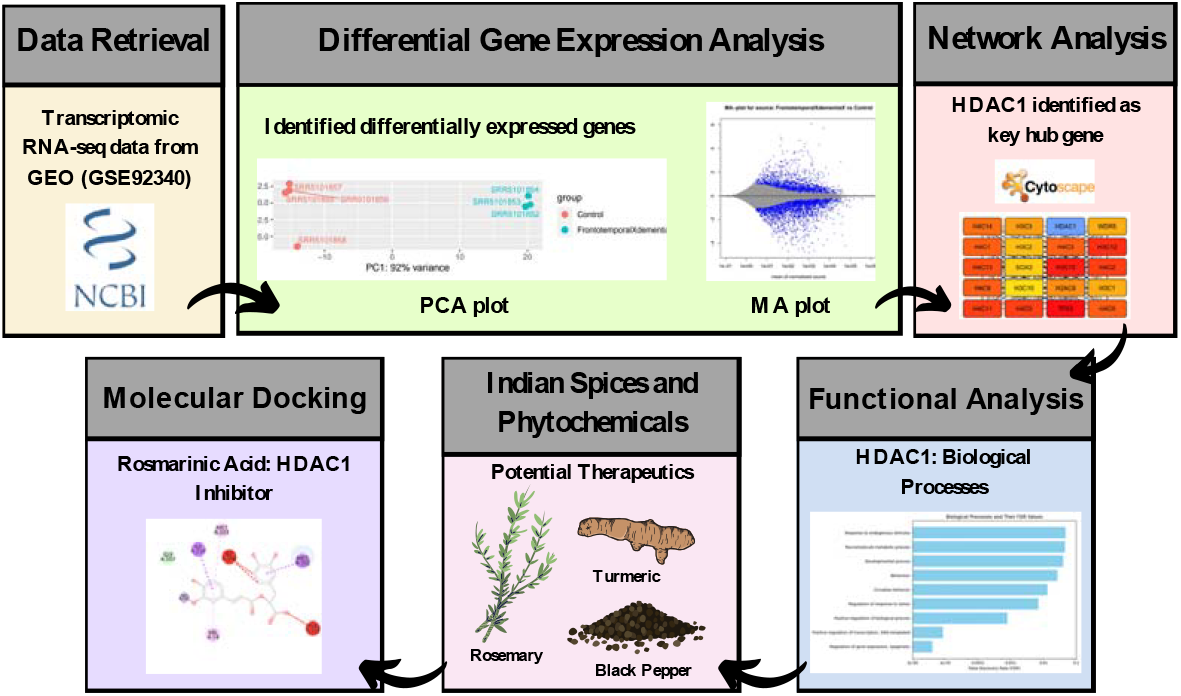
Flowchart illustrating the complete CADD pipeline.

### Data retrieval

This study highlights the importance of conducting transcriptomic sequence analysis on both patient and control samples. The primary platform used for data acquisition and gathering relevant information was the GEO [11] databases hosted by the National Centre for Biotechnology Information (NCBI).

### Quality control and trimming

The quality assessments for each sample were done using FASTQC [12]. FASTQC is a powerful tool used for examining the quality of raw sequence data derived from high-throughput sequencing data. To ensure the reliability of our data, we later used the Trim (Trimmomatic) tool [13] to eliminate low-quality reads and duplicated sequences from samples. For automated quality and adapter trimming, which includes the removal of biased methylation positions, we used the Trim tool. Both FASTQC and Trim were accessed through Galaxy Europe [14].

### Alignment and read counts

For aligning the clean reads to the *Homo sapiens* reference genome (GRCh38), we used HISAT-2 [15], a valuable tool available via Galaxy Server Europe. This alignment process is important as it accurately reads to their respective genetic locations, setting the stage for following gene expression analysis. To evaluate gene expression levels in our samples, we used a tool called FeatureCounts [16]. This tool enabled us to count the number of reads associated with each gene after the alignment process was completed.

### Identifying differentially expressed genes

Differentially expressed genes in the patient sample were analyzed using DESeq2 [17] (Differential Expression analysis of RNA-Seq data using the Negative Binomial distribution, Sequencing depth, and pair-wise models) DESeq2 is a dependable statistical method that accurately identifies genes displaying significant expression differences between distinct sample groups. We considered genes that are more active in FTD patients than in the control group using Padj and log2(FC) values. The criteria set for identifying upregulated genes were Padj values <0.05 and log2(FC) values >0.

### Gene Target Interaction Network Analysis and Functional Identification of Hub Genes

To explore the interactions and functions of the upregulated genes, we utilized the freely available Cytoscape software, which is specifically designed for analyzing complex biological networks. Additionally, we used the Cytoscape STRING plugin [18], limiting our analysis to protein-protein interactions with a confidence value of 0.70 or higher, as backed by dependable experimental evidence. To identify the most critical nodes in our network analysis, we used the CytoHubba plugin [19] within Cytoscape. This tool employs various topological techniques to determine and explore essential components and sub-networks within complex interactomes. CytoHubba uses six different importance metrics rooted in the concept of shortest paths, including Bottleneck (BN), Eccentricity, Closeness, Radiality, Betweenness, and Stress. Additionally, it employs distinct ranking methods such as Degree, Edge Percolated Component (EPC), Maximum Neighbourhood Component (MNC), Density of Maximum Neighbourhood Component (DMNC), and Maximal Clique Centrality (MCC) [19]. Comprehensive functional analysis to get deep insights into biological processes and pathways associated with the upregulated genes identified in our study. This crucial step involved the use of databases, including String Database, to understand cellular components, molecular functions, and biological processes in which these genes or proteins are involved. Our aim is to understand their roles through Gene ontology (GO) terms [20]. String Database (https://string-db.org/), this database compiles interactions from various sources, providing insights into Gene Ontology terms and improving our understanding of hub genes through Network Analysis. It provides useful information for biological process analysis.

### Assessing Medicinal Properties of Indian Spice Phytochemicals

Indian spices are rich in active phytochemicals. They are used for both culinary and medicinal purposes. These spices are known for their unique flavours and health benefits [21]. Some Indian spices contain active phytochemicals with antipsychotic properties. Antipsychotic agents can influence neurotransmitters and receptors involved in the treatment of psychosis-related conditions and symptoms [22]. Psychosis is often associated with conditions like dementia and other psychotic disorders [23]. Molinspiration (https://www.molinspiration.com/), is an online tool to measure medicinal properties of Indian spice phytochemicals. We gathered eighty-one antipsychotic phytochemicals. Molinspiration calculates key molecular properties like molecular weight, lipophilicity, hydrogen bond donors and acceptors, topological polar surface area, rotatable bonds, and Lipinski’s rule of five violations [24]. It predicts bioactivity scores for six drug target classes, indicating the likelihood of a molecule binding to a specific target based on its structural features. Increased bioactivity scores suggest a greater potential for effective binding.

### Predictive Molecular Properties and Bioactivity Scores

Predicting toxicity is necessary for identifying potential safety concerns and reducing the risk of adverse effects associated with pharmaceutical compounds. Toxicity testing is done using computational methods like ProTox-II (https://tox-new.charite.de/protox_II/), which predicts toxicity by calculating the properties of compounds based on their chemical structure. This method is cost-effective, fast, and ethical. ProTox-II is a web server that predicts toxicity outcomes for compounds using various approaches like molecular similarity, pharmacophore analysis, fragment propensities, and machine-learning models. It generates comprehensive toxicity predictions for compounds across thirty-three different models. Recently, we used ProTox-II to evaluate the toxicity of twenty-six phytochemicals occurring naturally in Indian spices. We established specific limits for selecting safe and drug-like phytochemicals, including compliance with Lipinski’s rule of five [24] and examining the overall Predicted Toxicity Class.

### Exploring HDAC1 Inhibition through Molecular Docking Analysis

Molecular docking is a computational simulation technique that can be used to identify potential inhibitors of target molecules [25, 26] (HDAC1), a protein that plays a critical role in regulating gene expression. However, currently available HDAC1 inhibitors lack specificity, meaning they can also inhibit other proteins in addition to HDAC1. This lack of specificity can lead to unwanted side effects and toxicity. In addition, many of the existing HDAC1 inhibitors are highly toxic, which limits their potential use as therapeutic agents. Our approach employs PatchDock (https://www.cs.tau.ac.il//~ppdock/PatchDock/) [10], the web server uses shape complementarity to enable molecular docking for exploring the pharmacological properties of active compounds found in Indian spices. The approach identifies surfaces of proteins and ligands, aligning them according to shape and electrostatic traits. Docking analyses were conducted for each phytochemical, enabling the assessment of their binding affinity and interactions based on selection criteria such as geometric shape complementarity score, atomic contact energy, hydrogen bonds, and clashes.

## Result and discussion

### Data Retrieval

Conducting extensive research on transcriptomic sequencing in patients with Frontotemporal Dementia, using the GEO database of NCBI to determine and categorize relevant studies. Using the GEO Accession viewer, we identified key data from document number GSE92340, which utilized RNA sequencing to investigate variations in gene expression in neurons from FTD patients. Our study included four control samples and three patient samples with cortical neuron iPSC (Fibroblast) cell/tissue type. These crucial steps allowed us to comprehensively analyze the molecular landscape of FTD.

### Identifying differentially expressed genes

PCA (Principal Component Analysis) plot shows distributing samples based on their variance along two principal components PC1 and PC2. The samples are grouped into two categories control and frontotemporal dementia. Through plot shown in Figure 2, samples are Clustered or separated on basis of their variance along these principal components, with PC1 having a much larger impact on variance than PC2. DESeq2 examined differential gene expression. Significantly expressed genes with positive log2FoldChange were observed to be upregulated that is 1882 and negative ones were observed to be downregulated that is 2747. The gene that was identified as upregulated, that is, had a p-value less than 0.05, and a log2foldchange value positive was taken into consideration for further analysis. MA plot highlights differentially expressed features between frontotemporal dementia and control group. The features that are upregulated in frontotemporal dementia are marked in blue in the plot shown in Figure 3.

**Figure 2:**
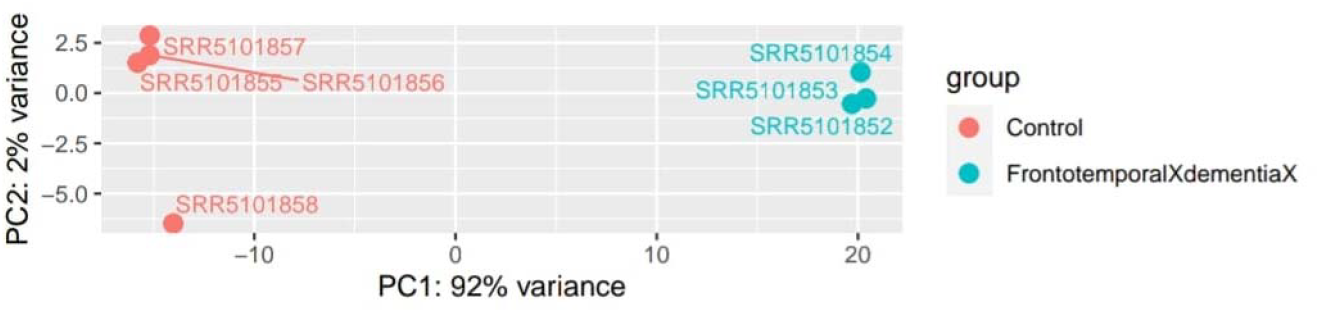
Principal component analysis (PC A) plot evaluating the variance between the samples.

**Figure 3:**
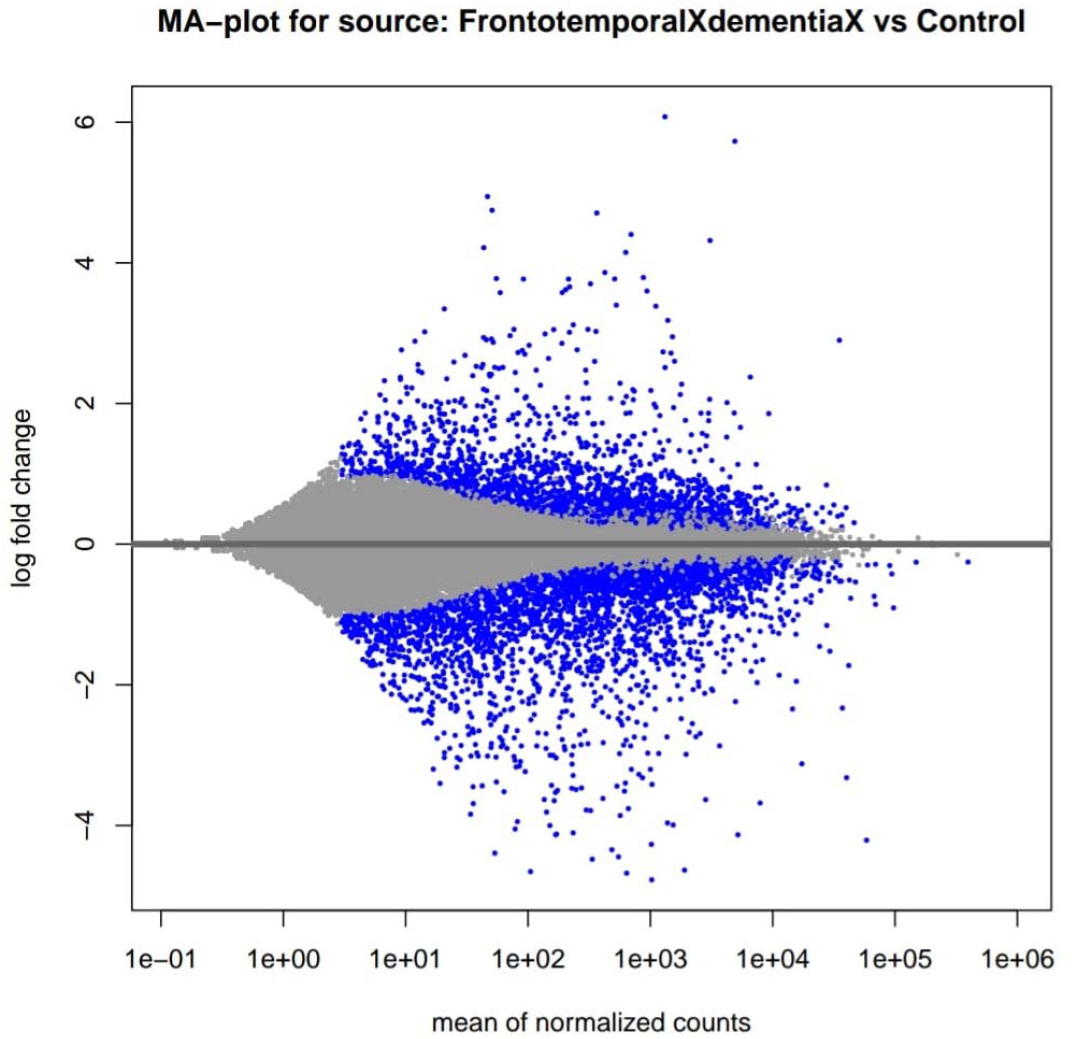
MA plot highlights differentially expressed features between Frontotemporal Dementia and Control groups.

### Network Analysis and Identification of Hub Genes

Using the STRING plugin in Cytoscape software, a gene-gene interaction network was constructed by utilising all upregulated genes. The network was filtered to a high confidence level of 0.70 and resulted in 1508 nodes and 2856 edges from the original 1882 genes. This comprehensive network provided valuable insights into complex chemical interactions and potential functional relationships among the upregulated genes studied. Please refer to Fig 2 for a visualisation of the interaction network created in Cytoscape. Through our analysis of various topological algorithms, it has become clear that HDAC1 is a key hub gene in our network. As a histone deacetylase, HDAC1 plays a vital role in regulating chromatin structure and gene expression. With a high degree score of 80, indicating numerous connections to other nodes in the network, HDAC1 also scored highly in other assessments such as Stress, Closeness, and Betweenness. These findings demonstrate the importance of HDAC1 in our network analysis and suggest its potential as a key factor in the molecular landscape. Please refer to Table 1 for further information on the parameter and score of the HDAC1 gene as obtained through Cytoscape’s CytoHubba Plugin.

**Table 1:**
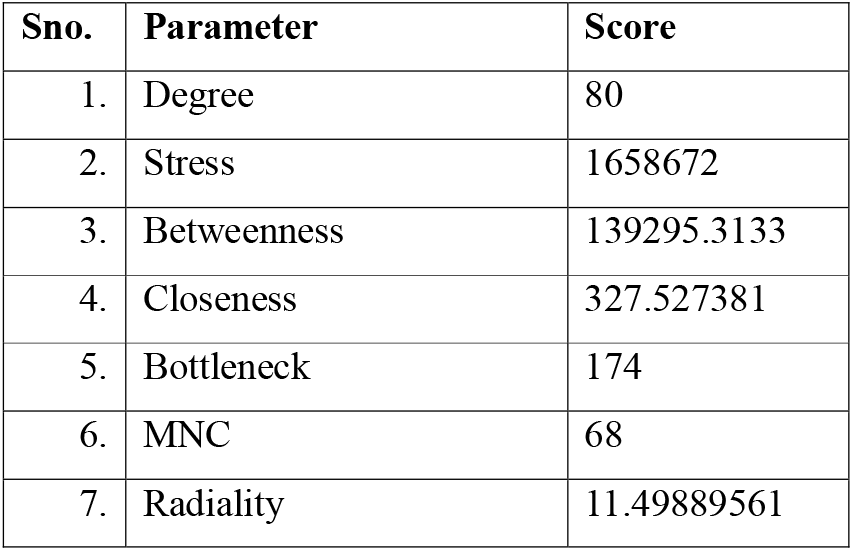
Displays the Parameter and score of the HDAC1 gene obtained from Cytoscape’s CytoHubba Plugin.

### Functional Analysis of Hub Gene

Hub gene HDAC1 appeared as a highly connected gene in our analysis. It codes key proteins that are involved in modifying chromatin structure. HDAC1 affects gene expression by modifying histones, which package DNA. This limits access to transcription factors needed for gene regulation chromatin compaction and access to the transcription factors required for gene regulation. We found that the HDAC1 gene is involved in important biological processes using the String Database. On Gene Ontology Analysis, it was observed that target genes HDAC1 play an essential role in important biological processes such as positive regulation of transcription (DNA-templated), behaviour, and Regulation of stress response as shown in Figure 4 Biological Processes with their False Discovery Rate Values (FDR).

**Figure 4:**
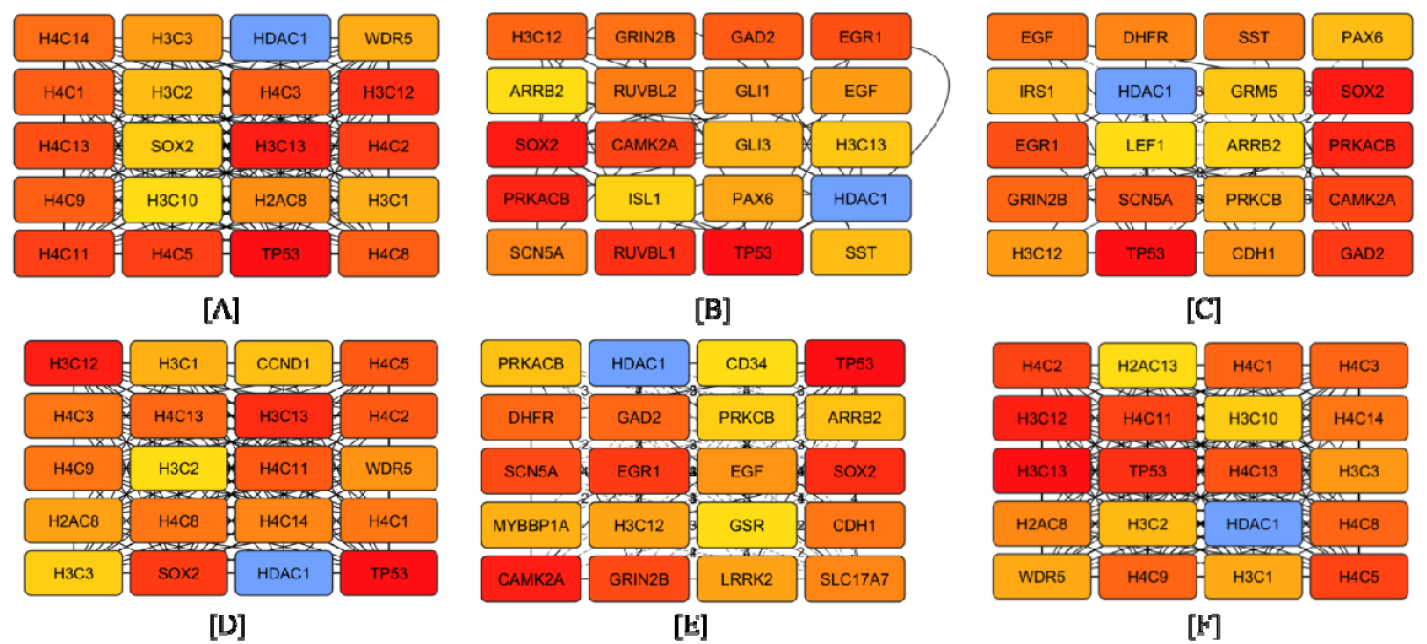
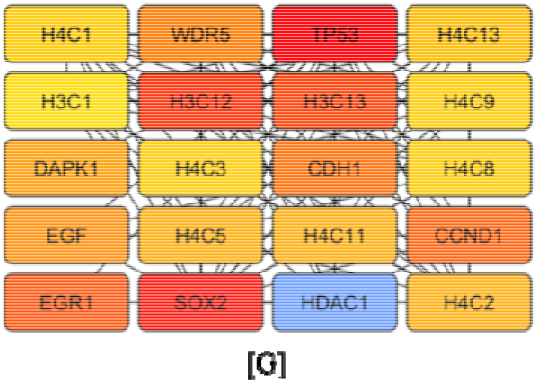
Network of hub genes in different parameter obtained from Cytoscape’s CytoHubba Plugin **[A]:** *Network of hub genes in top 20 for Degree parameter obtained from Cytoscape’s CytoHubba Plugin*, **[B]:** *Network of hub genes in top 20 for Stress parameter obtained from Cytoscape’s CytoHubba Plugin*, **[C]:** *Network of hub genes in top 20 for Betweenness parameter obtained from Cytoscape’s CytoHubba Plugin*, **[D]:** *Network of hub genes in top 20 for Closeness parameter obtained from Cytoscape’s CytoHubba Plugin*, **[E]:** *Network of hub genes in top 20 for Bottleneck parameter obtained from Cytoscape’s CytoHubba Plugin*, **[F]:** *Network of hub genes in top 20 for MNC parameter obtained from Cytoscape’s CytoHubba Plugin*, **[G]:** *Network of hub genes in top 20 for Radiality parameter obtained from Cytoscape’s CytoHubba Plugin*

**Figure 5:**
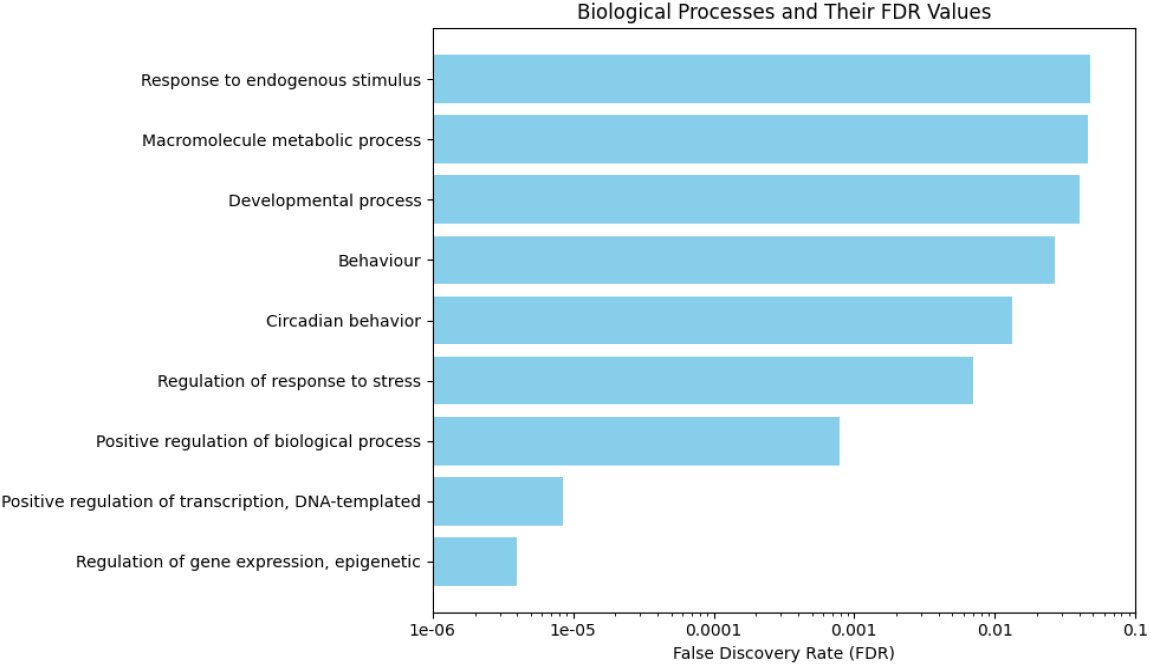
Bar graph for Biological Processes with False Discovery Rate Values (FDR).

### Assessing Medicinal Properties of Indian Spice Phytochemicals

Molinspiration identifies essential molecular properties and predicts bioactivity scores against different drug targets. 81 Indian spices were searched that have Antipsychotic properties, and twenty-six active phytochemicals were found that follow Lipinski’s rule, which is a guide for drug-likeness. These findings provide safer and more effective alternatives for treating psychosis-related conditions. Table 2 shows the list of Indian Spices and Their Active Phytochemicals with their references.

**Table 2:** Indian Spices and Their Active Phytochemicals with references.

| Sno. | Indian Spices | Scientific Name | Active Phytochemical | Ref |
| --- | --- | --- | --- | --- |
| 1. | Rosemary (Rusmari) | *Rosmarinus officinalis L* | Rosmarinic acid | [27, 28] |
| 2. | Turmeric (Haldi) | *Curcuma longa* | Curcumin | [29] |
| 3. | Black pepper (Kali mirch) | *Piper guineense* | Piperine | [30] |
| 4. | Long pepper (Pippali) | *Piper longum* | Piperlongumine | [31] |
| 5. | Ginger (Adrak) | *Zingiber officinale* | Gingerol | [32] |
| 6. | Celery seeds (Ajmoda) | *Apium graveolens* | Apigenin | [33] |
| 7. | Red sandalwood<br>(Raktachandan) | *Pterocarpus santalinusn*<br>*L.* | Santalin | [34] |

### Predictive Molecular Properties and Bioactivity Score

ProTox-II analysis showed that twenty-three out of twenty-six phytochemicals are safe and have drug-like features. This discovery is promising for identifying potential therapeutic agents and developing safer, more effective drugs. Table 3 and Figure shows a list of 23

**Table 3:**
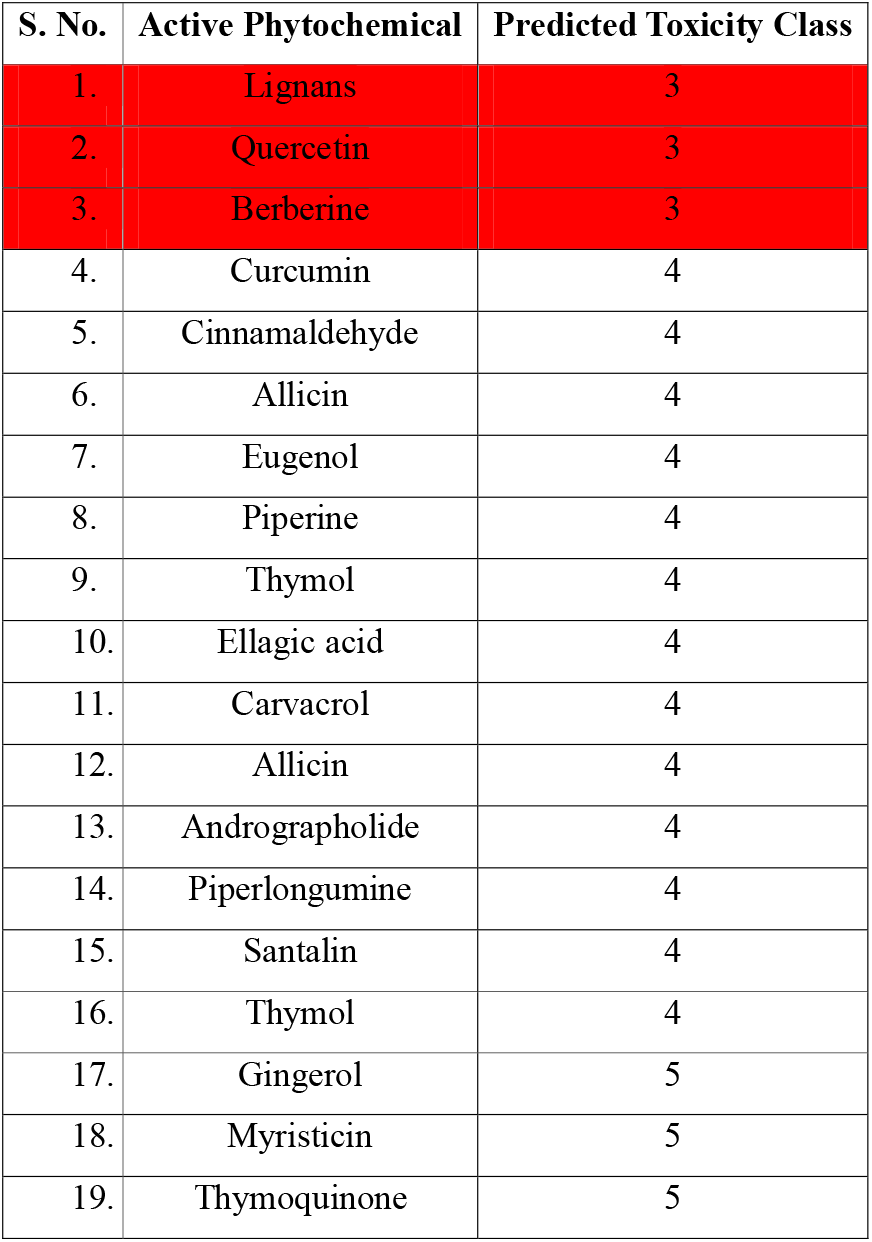
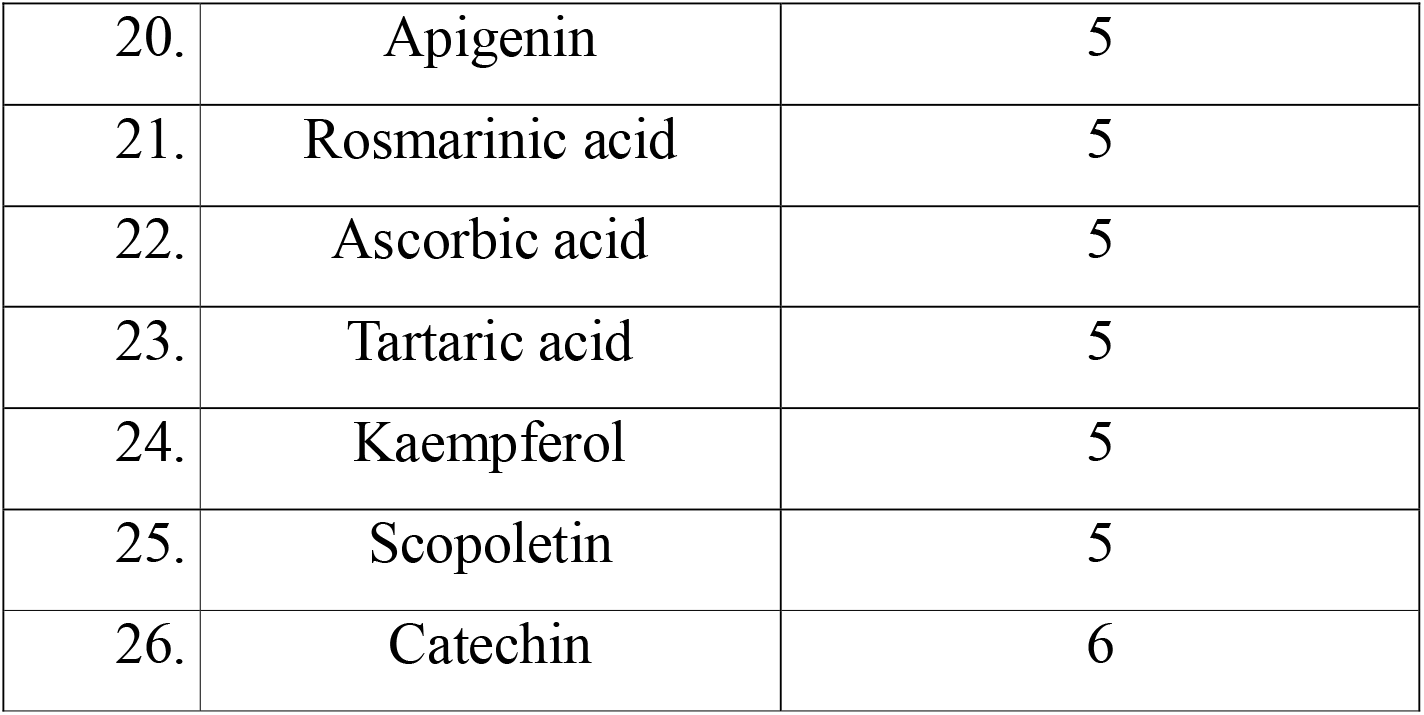
List of 23 Active Phytochemicals with Predicted Toxicity Class using ProToxII. Active phytochemicals with predicted Toxicity Classes using ProToxII.

| S. No. | Active Phytochemical | Predicted Toxicity Class |
| --- | --- | --- |
| 1. | Lignans | 3 |
| 2. | Quercetin | 3 |
| 3. | Berberine | 3 |
| 4. | Curcumin | 4 |
| 5. | Cinnamaldehyde | 4 |
| 6. | Allicin | 4 |
| 7. | Eugenol | 4 |
| 8. | Piperine | 4 |
| 9. | Thymol | 4 |
| 10. | Ellagic acid | 4 |
| 11. | Carvacrol | 4 |
| 12. | Allicin | 4 |
| 13. | Andrographolide | 4 |
| 14. | Piperlongumine | 4 |
| 15. | Santalin | 4 |
| 16. | Thymol | 4 |
| 17. | Gingerol | 5 |
| 18. | Myristicin | 5 |
| 19. | Thymoquinone | 5 |
| **Exploring Inhibition Molecular Analysis** |  |  |
| 20. | Apigenin | 5 |
| 21. | Rosmarinic acid | 5 |
| 22. | Ascorbic acid | 5 |
| 23. | Tartaric acid | 5 |
| 24. | Kaempferol | 5 |
| 25. | Scopoletin | 5 |
| 26. | Catechin | 6 |

### Exploring HDAC1 Inhibition through Molecular Docking Analysis

Rosmarinic acid isolated from Rosemary (*Rosmarinus officinalis L*) highly suggests HDAC1 gene inhibitor in our molecular docking analysis. Rosmarinic acid shows potential as therapeutic mediation for Frontotemporal Dementia. It can prevent the harmful effects of HDAC1 gene dysregulation and protein aggregation. Molecular docking analysis was used in our study to assess its ability, we examined the interaction between the HDAC1 protein and Rosmarinic acid an Active phytochemical found in Rosemary. Our analysis showed that rosmarinic acid binds strongly to the HDAC1 binding pocket as shown in Figure 6, suggesting it could be a promising therapeutic agent for Frontotemporal Dementia. Rosmarinic acid shows potential as an HDAC1 gene inhibitor or modulator, according to PatchDock results as shown in Table 4. The interaction has a high score, significant interface area, favourable high ACE value, and specific spatial coordinates as shown in Figure 7. Further analysis is recommended for therapeutic implications.

**Table 4:** Indian Spices with their Active Phytochemical compound having ACE values above −300 kcal/mol.

| Sno. | Indian Spices | Scientific Name | Active Phytochemical | ACE |
| --- | --- | --- | --- | --- |
| 1. | Rosemary (Rusmari) | *Rosmarinus* | Rosmarinic acid | -395.48 |

|  |  | *officinalis L* |  |  |
| --- | --- | --- | --- | --- |
| 2. | Turmeric (Haldi) | *Curcuma longa* | Curcumin | -387.93 |
| 3. | Black pepper (Kali mirch) | *Piper guineense* | Piperine | -328.92 |
| 4. | Long pepper (Pippali) | *Piper longum* | Piperlongumine | -326.53 |
| 5. | Ginger (Adrak) | *Zingiber officinale* | Gingerol | -318.32 |
| 6. | Celery seeds (Ajmoda) | *Apium graveolens* | Apigenin | -314.2 |
| 7. | Red sandalwood<br>(Raktachandan) | *Pterocarpus*<br>*santalinusn L.* | Santalin | -306.55 |

**Figure 6:**
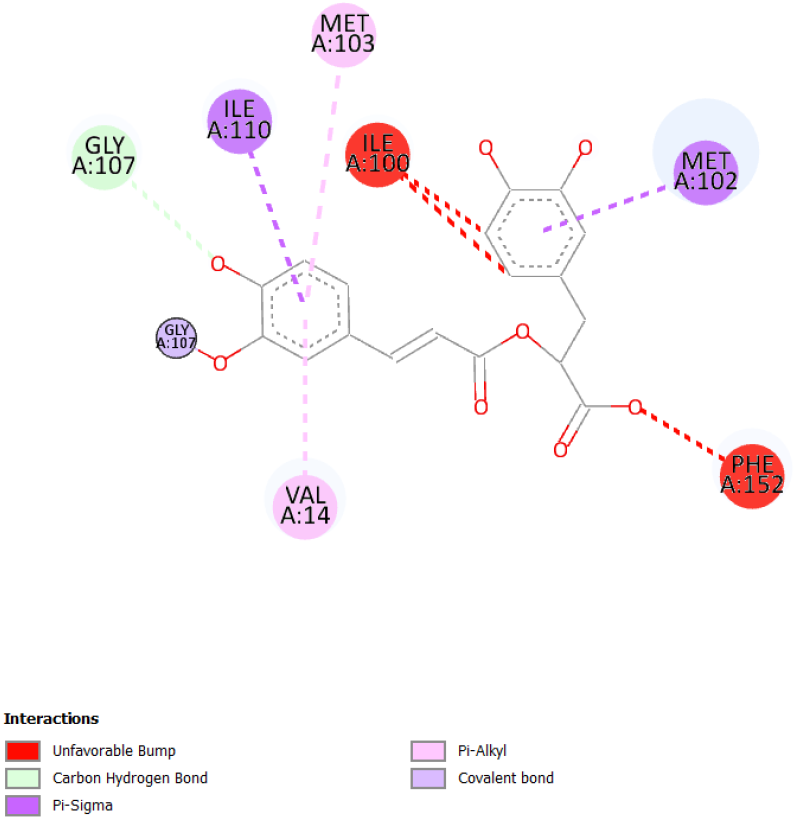
2D Structure of Receptor-Ligand Interaction using Discovery Studio Visualizer

**Figure 7:**
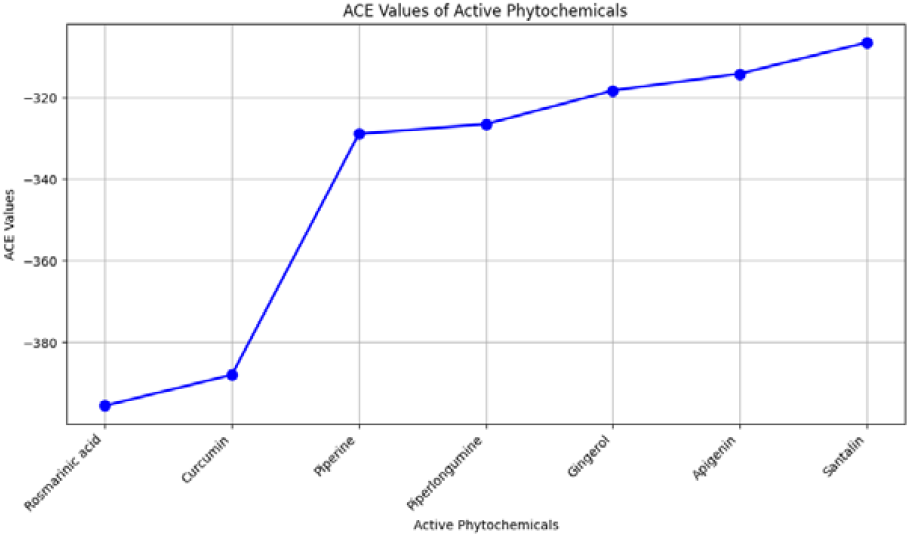
ACE value with Active Phytochemicals.

## Conclusion

The study conducted an extensive analysis to understand the complex landscape of frontotemporal dementia (FTD) [1] using a multifaceted approach that involved transcriptomic analysis, network exploration, functional annotation, and innovative screening of phytochemical compounds. To begin with, Galaxy Server Europe [15] was utilized to identify differentially expressed genes (DEGs) [5] that are associated with FTD, which led to the discovery of significant molecular alterations compared to healthy controls. The subsequent network analysis, performed using Cytoscape [6] and STRING database [7], unravelled complex gene-gene interactions, revealing HDAC1 as a pivotal hub gene that plays vital role in chromatin modulation [35], gene expression regulation [36], behavioural affects [37] and epigenetic regulation [38]. Overall, the study employed innovative techniques and tools to gain deeper understanding of molecular mechanisms involved in FTD, which could help in the development of effective therapeutic strategies to treat this debilitating disease.

Functional analysis has been beneficial in revealing the involvement of HDAC1 in certain of the most crucial biological processes such as transcriptional regulation [39] and stress response [40]. This has led to the crucial role played by HDAC1 in the pathophysiology of FTD (Frontotemporal Dementia). Additionally, detailed analysis of phytochemicals found in Indian spices has led to the discovery of promising candidates that possess antipsychotic properties. These candidates have been identified to possess drug-like features and safe profiles as assessed through Molinspiration and ProTox-II [9] analyses. This information can be particularly beneficial in advancing research on FTD and in development of new drugs with antipsychotic properties.

It is quite remarkable that Rosmarinic acid, natural compound found in Rosemary [27], has been identified as a potent inhibitor of HDAC1 (histone deacetylase 1) [37]. The compound has demonstrated significant binding affinity towards HDAC1, and this finding has potential therapeutic implications for frontotemporal dementia (FTD). Molecular docking analyses conducted via PatchDock [10] have revealed that Rosmarinic acid is capable of interacting strongly with HDAC1. This suggests that it can be used as viable therapeutic agent against FTD by mitigating HDAC1 dysregulation and protein aggregation. These promising results lead to further research and development of Rosmarinic acid as a potential treatment for FTD. This extensive study focuses on the complex molecular mechanisms that contribute to frontotemporal dementia (FTD), a neurodegenerative disease that affects cognitive and behavioural functions. The research sheds light on potential biomarkers and therapeutic targets that could be crucial in developing effective treatments for this condition. Rosmarinic acid, a natural compound found in herbs such as rosemary and sage, is a promising therapeutic mediator that emerged from the study. The findings suggest that Rosmarinic acid could be effective in reducing the effects of FTD and improving patients’ quality of life. In addition, these new insights provide exciting new avenues for research and intervention in the field of frontotemporal dementia.

